# Quantifying the prevalence of assortative mating in a human population

**DOI:** 10.1101/848911

**Authors:** Klaus Jaffe

## Abstract

For the first time, empirical evidence allowed to construct the frequency distribution of a genetic relatedness index between the parents of about half a million individuals living in the UK. The results suggest that over 30% of the population is the product of parents mating assortatively. The rest is probably the offspring of parents matching the genetic composition of their partners randomly. High degrees of genetic relatedness between parents, *i.e.* extreme inbreeding, was rare. This result shows that assortative mating is likely to be highly prevalent in human populations. Thus, assuming only random mating among humans, as widely done in ecology and population genetic studies, is not an appropriate approximation to reality. The existence of assortative mating has to be accounted for. The results suggest the conclusion that both, assortative and random mating, are evolutionary stable strategies. This improved insight allows to better understand complex evolutionary phenomena, such as the emergence and maintenance of sex, the speed of adaptation, runaway adaptation, maintenance of cooperation, and many others in human and animal populations.

## INTRODUCTION

In most population genetic models, random mating is considered the default option. Assortation, rather than random mating, has been proposed to be necessary and widespread in sexual reproduction of most living beings (8). Assortative mating has been demonstrated to occur among humans (1,5,14) and many other animals and plants (7). The physical-dynamic mechanism behind this assumption is that assortation allows evolution to handle epistasis more effectively than random mating, increasing the error threshold for lethal mutations (11,12), allowing synergy between cooperating genes to be maintained and expanded (9). Assortative mating is also known to accelerate adaptation due to its effect on the Hardy–Weinberg equilibrium (13). As these mechanism are basic features of all complex evolutionary scenarios, this has implications for a wide range of disciplines (6).

The degree to which non-random mating influences the genetic architecture remains unclear as most theoretical studies assume random mating. However, existing simulations showed that assuming non-random mating has a substantial effect on the outcomes (2,10). Empirical research (14) studied genetic variants associated with human height to assess the degree of height-related assortative mating in European-American and African-American populations. The study compared the inbreeding coefficient estimated using known height associated variants with that calculated from frequency matched sets of random variants. The results showed a significantly higher inbreeding coefficients for the height associated variants than from frequency matched random variants (P < 0.05), demonstrating height-related assortative mating in both populations.

Recent advances in quantitative genetics allow to access empirical information about the prevalence of mate selection mechanisms acting on a large set of genetic traits in natural populations. Specifically, data in the UK Biobank allowed to construct the frequency distribution of genetic relatedness of the parents of about half a million people living in Britain. Thus, for the first time, empirical evidence from large human populations, as to the generality of the different mating strategies (random or assortative), is available (4). Here I use this data to assess the extent to which assortative mating, assessed through genomic means, occurs in a human population.

## METHODS

I used data gathered for another purpose (15-16) but which has the information required to build frequency distributions of the genetic relatedness between parents (4). The form of this frequency distribution allows to discern which of the two mate selection mechanisms, random or assortative, is more prevalent. Random mating should produce an inbreeding coefficients frequency distribution among individuals of a population that matches a truncated Poisson distribution with the maximum close to the mean of the inbreeding coefficients of that population. Out-breeding should produce a frequency distribution where low inbreeding coefficients are more common. Assortative mating should favor higher inbreeding coefficients, than those expected from populations with only random mating, producing a skewed distribution with a fat tail towards high genetic relatedness. Extreme inbreeding is known to depress the fitness of the affected individuals (15) and should thus be rare.

Genotyped Single Nucleotide Polymorphism (SNPs) from Biobank were used to detect large runs of homozygosity (ROH). Runs of Homozygosity (ROH) are genomic regions where identical haplotypes are inherited from each parent. Once ROHs was detected, an inbreeding measure Froh was calculate for each individual as a measure of autozygosis by dividing the cumulated length of ROH in Mb by an estimate of the length of the human autosome (15).

The percentage of individuals born through assortative mating will affect the form of the frequency distribution of F. A theoretical distribution of the degree of assortment assuming 100% random mating was produced using Monte Carlo matching of two randomly chosen haploid DNA strands in a virtual population of 500 individuals each possessing 100 loci containing one of potentially 50 different alleles. The model (see supporting material) was a simplified version of virtual genomes where ROHs were represented by virtual alleles. Autozygosis was estimated in the simulation through the number of homologous allelic matches producing the inbreeding estimate “Frnd”. The frequency distribution of Frnd was normalized to the number of individuals in the sample (456414).

Statistics is often more misleading than useful (17,18). Comparing the randomly generated results with the empirically obtained frequency distribution produces probability values of rejecting the null hypothesis close to absolute 0. This is misleading because many other uncertainties, not captured by this statistics, affect the comparison. For example, the structure of allele combinations and their distribution in the genome affects this result, and so do other factors, discussed below. Thus, I will avoid impressing the reader with astronomically small probabilities in order to focus on the information that can humbly and rationally be extracted from the empirical data presented.

## RESULTS

Two frequency distributions of genetic relatedness of parents are presented. The Frnd distribution of random matches of identical alleles obtained with a Mont Carlo model, mimicking the outcome if all parents of the individuals engaged in random mating; and the Froh distribution of the inbreeding coefficient obtained from data in Biobank. Both are plotted in Figure 1a and 1b. Clearly, the effect of assortation of the frequency distribution of Froh is evident when contrasting it with the distribution of Frnd simulating random mating.

**Figure 1.**
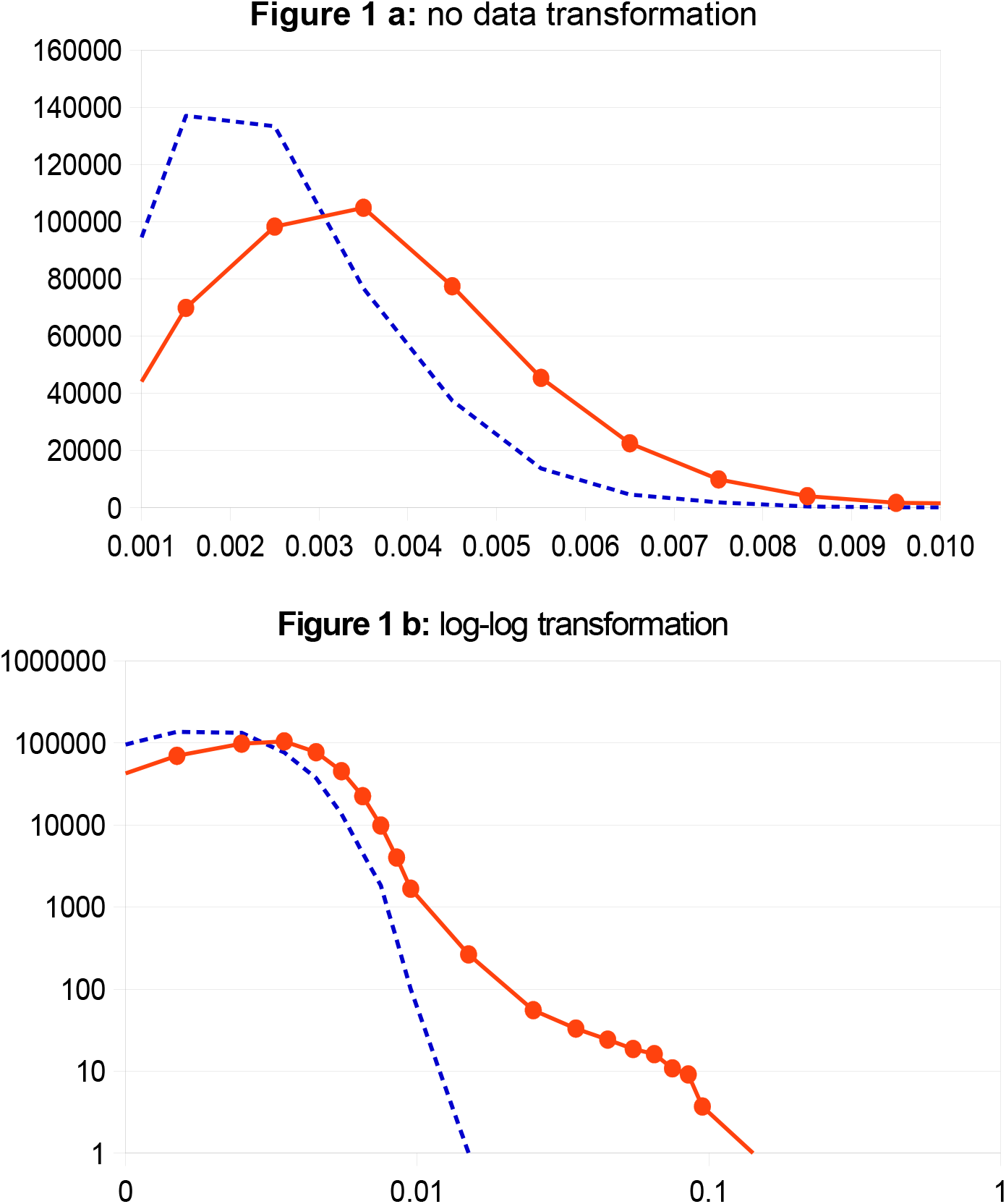
Percentage of the number of individuals from the Biobank database at F value intervals of 0.001 Froh units used as an inbreeding coefficient. The red continuous line represents the actual experimental data, the blue punctuated line are estimates for Frnd from simulations assuming random mating.

The data shows that the average value of Froh is 0.0036, the maximum value in the Froh curve ocurrs at values of 0.0025, and the median is above 0.01 Froh; indicating that the curve is very skewed towards the right of the plot, i.e. towards higher values of Froh. Compared to an expected normal Poisson distribution. Also, the fat tail on the right of the Froh curve indicate that the amount of individuals with a higher Froh than the median is substantial. The figure shows that for values of F > 0.01, the frequency of individuals diminishes markedly in the Biobank data, congruent with finding that extreme inbreeding is non-adaptive as shown elsewhere (3, 15).

The left part of the curve is truncated due to the nature of the empirical data. The number of alleles studies is still small in Biobank. The parameters of the theoretical simulations were adjusted so as to mimic this handicap in the empirical data. This limitation does not allow to infer a putative prevalence of dissortative mating or extreme outbreeding.

## DISCUSSION

A structural problem in comparing empirical and theoretical data is the matching of parameter values. In our case, the matching between Froh and Frnd. For more robust conclusions, independent evidence obtained by other means is required (see below). Figure 1b minimize the effect of mismatch errors by presenting a log-log plot. The result shows that the inbreeding coefficient from Biobank data between the parents of the individuals studied, produced a maximum peak at Froh’s much higher than the mean value of the population. This value is also far higher than that estimated by numerical simulations assuming random mating.

Natural human populations have, important genetic homologies due to the fact that they configure a single species with a common ancestor. This fact is not captured by the random simulation model, as data from SNP was also chosen from parts of the genome showing important variance. But clearly, the frequency distribution of actual empirical F data is very different from the simulations which show no fat tail. In Figure 1, 30% of the population has higher F scores than predicted by simulations of random mating. This proportion is probably higher as not all genes relevant to assortation were included in this study. The evidence hints then that assortative mating is an adaptive behavior. This means that ignoring assortative mating, as is common in population genetic studies, is not a rational choice.

Assortative mating can be achieved by passive and active means. Geographic isolation of sub-populations (passive assortment) can achieve similar levels of assortment as that of active mate selection. If assortment is an adaptive feature of humans, however, individuals whose parents engaged in assortative mating will show higher F values and should have a higher fitness. A proof that assortation is at work can be provided by comparing a fitness index for extreme out-breeders (Froh 0.001) with that of random maters (Froh 0.004), with those mating assortatively (Froh 0.02-0.03) and with extreme in-breeders (Froh > 0.1). This last group was shown to have phenotypic fitness means between 0.3 and 0.7 standard deviation below the population mean for 7 traits, including stature and cognitive ability, consistent with inbreeding depression estimated from individuals with low levels of inbreeding (1). The falsifiable prediction is that offspring from assortative maters should have fitness values above the population mean. Science has now the tools to study the effect of homozygosis to understand genetic epistatic synergies favored by assortation. This new focus allows a much better understanding of complex phenomena, such as the emergence and maintenance of sex among creatures with complex genomes (i.e. all living beings).

This study was motivated by computer simulations predicting the prevalence of assortative mating among sexual beings (8). The empirical results presented here close a loop of the cyclic interaction between theory and empirical results. Novel knowledge was unveiled from this dynamic interaction: both, random mating and assortative mating occur in human populations, suggesting that both are evolutionary stable strategies. Passive and active assortative mating synergize favoring speciation (19). Therefore, random mating and dissortative mating must exist among members of a single species that is not splitting. The balance between these strategies, however, might vary in space and time. An optimum balance between them might exist. Further interactions between theory and empirical evidence should help us unveil these questions.

## Acknowledgment

I thank Peter Visher and Loic Yengo Dimbou for access to the data empirical presented here

## Notes

https://github.com/loic-yengo/Histogram-of-inbreeding-coefficients-in-the-UKB

